# Laminar pattern of adolescent development changes in working memory neuronal activity

**DOI:** 10.1101/2023.07.28.550982

**Authors:** Junda Zhu, Benjamin M. Hammond, Xin Maizie Zhou, Christos Constantinidis

**Author notes:** Address correspondence to: Christos Constantinidis, PhD.

## Abstract

Adolescent development is characterized by an improvement in cognitive abilities, such as working memory. Neurophysiological recordings in a non-human primate model of adolescence have revealed changes in neural activity that mirror improvement in behavior, including higher firing rate during the delay intervals of working memory tasks. The laminar distribution of these changes is unknown. By some accounts, persistent activity is more pronounced in superficial layers, so we sought to determine whether changes are most pronounced there. We therefore analyzed neurophysiological recordings from neurons recorded in the young and adult stage, at different cortical depths. Superficial layers exhibited increased baseline firing rate in the adult stage. Unexpectedly, changes in persistent activity were most pronounced in the middle layers. Finally, improved discriminability of stimulus location was most evident in the deeper layers. These results reveal the laminar pattern of neural activity maturation that is associated with cognitive improvement.

**NEW AND NOTEWORTHY:** Structural brain changes are evident during adolescent development particularly in the cortical thickness of the prefrontal cortex, at a time when working memory ability increases markedly. The depth distribution of neurophysiological changes during adolescence is not known. Here we show that neurophysiological changes are not confined to superficial layers, which have most often been implicated in the maintenance of working memory. Contrary to expectations, greatest changes were evident in intermediate layers of the prefrontal cortex.

## INTRODUCTION

Working memory, the ability to maintain and manipulate information in mind over a timescale of seconds, improves during the course of adolescence (1-4). Anatomical and functional changes in a network of cortical areas including the dorsolateral prefrontal cortex take place during this time and are predictive of performance gains in working memory tasks (5-10). Most importantly, cortical volume and thickness of the frontal lobe change considerably during adolescent development, with an overall pattern of decreased thickness evident in adulthood, thought to represent pruning of unwanted synapses (11, 12). Neurodevelopmental disorders that manifest themselves in late adolescence, most notably schizophrenia, are characterized by deficits of working memory and corresponding aberrations in dorsolateral prefrontal cortex function and cortical thickness (13-16).

Over the past few years, it has become possible to study the development and maturation of neuronal responses in the prefrontal cortex of non-human primates between the time of adolescence and adulthood (17, 18). Male rhesus monkeys (*Macaca mulatta*) enter puberty at approximately 3.5 years of age and reach full sexual maturity at age 5, aging roughly 3 times faster than humans (19, 20). Behavioral and neurophysiological recordings obtained two years apart span the greatest part of adolescence and have revealed substantial differences in activity and response properties of neurons between the adolescent and adult prefrontal cortex (21, 22). Most importantly, these include higher persistent firing rate in the delay periods of working memory tasks (21).

The laminar pattern of neurophysiological response changes has not been investigated until now. Strong anatomical (23, 24) and physiological evidence (25) suggests that superficial layers of the prefrontal cortex play a critical role in the generation of persistent activity and maintenance of working memory. Many previous studies of working memory have focused predominantly on superficial layers for this reason (26). However, there has also been evidence that deep layer neurons contribute to this activity (27). We were therefore motivated to examine neuronal responses obtained at different cortical depths and investigate the laminar maturation of the prefrontal cortex between developmental stages in a non-human primate model of working memory.

## MATERIALS AND METHODS

Four male rhesus monkeys (Macaca mulatta) were used in this study. All surgical and animal use procedures were reviewed and approved by the Wake Forest University Institutional Animal Care and Use Committee, in accordance with the U.S. Public Health Service Policy on humane care and use of laboratory animals and the National Research Council’s Guide for the care and use of laboratory animals.

### Developmental profiles

We tracked developmental measures of monkeys on a quarterly basis before, during, and after neurophysiological recordings, as we have documented in detail previously (21, 22). We obtained morphometric measures including body weight, crown-to-rump length, chest circumference, ulna and femur length, testicular volume, and eruption of canines. We additionally measured bone maturation by X-rays of the upper and lower extremities; and assayed serum concentration of circulating hormones including testosterone [T] and dihydrotestosterone [DHT]. Using these measures, we determined puberty onset, and behavioral and neurophysiological recordings were collected at two stages: mid-adolescence (‘young stage’) and adulthood (‘adult stage’), with a 1.6-2.1-year gap. Median ages at the mid-adolescent and adulthood stage were 4.3 years and 6.3 years, respectively.

### Behavioral Tasks

The monkeys were trained to perform an Oculomotor Delayed Response (ODR) task during the young stage (Fig. 1). Once the animals had reached asymptotic performance, neuronal recordings were obtained (described in the next section). At the conclusion of these recordings, the animals were returned to their colony and were no longer tested or trained in any task for a period of ∼1 year. A new period of testing and recording was performed in the adult stage.

**Figure 1.**
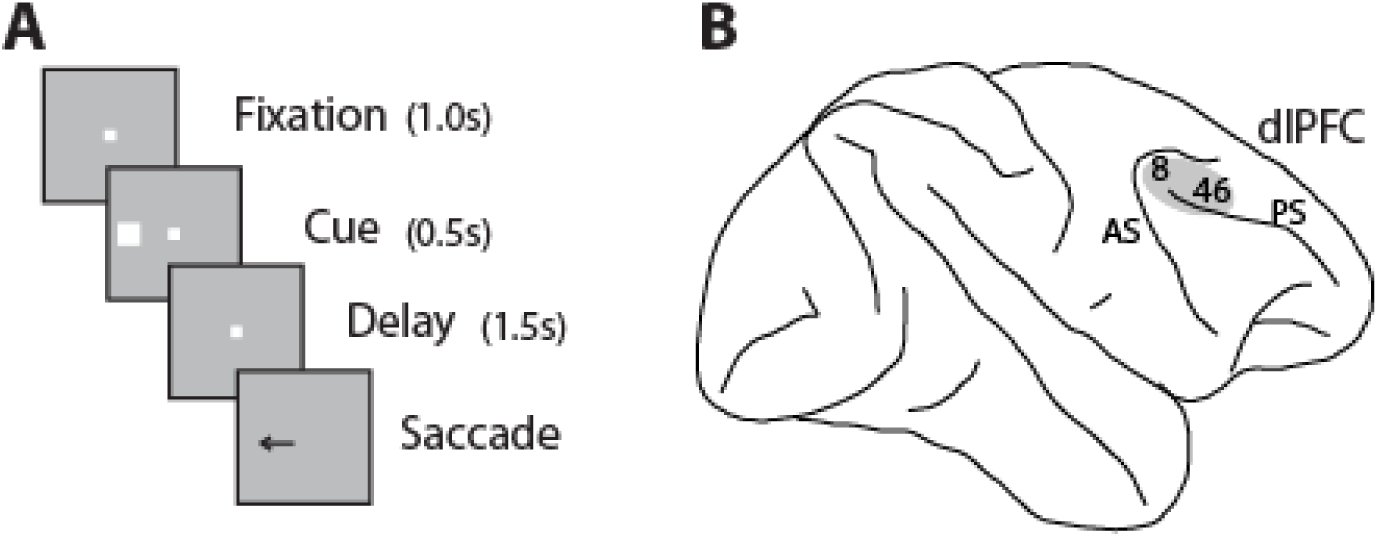
**A** Sequence of events in the Oculomotor Delayed Response (ODR) task. **B**. Schematic diagram of the monkey brain with approximate location of neuronal recordings in areas 8 and 46 of the dorsolateral prefrontal cortex (dlPFC) highlighted. Abbreviations: AS, arcuate sulcus; PS, principal sulcus.

The ODR task is a spatial working memory task requiring subjects to remember the location of a cue stimulus flashed on a screen for 0.5 s. In this study, the cue was a 1° white square stimulus that could appear at one of eight locations arranged on a circle of 10° eccentricity. After a 1.5 s delay period, the fixation point was extinguished, and the monkey was trained to make an eye movement to the remembered location of the cue within 0.6 s. The saccade needed to terminate on a 5–6° radius window centered on the stimulus (within 3–4° from the edge of the stimulus), and the monkey was required to hold fixation within this window for 0.1 s.

Animals were rewarded with fruit juice for successful completion of a trial. Eye position was monitored with an infrared eye tracking system (ISCAN, RK-716; ISCAN, Burlington, MA). Breaking fixation at any point before the offset of the fixation point aborted the trial and resulted in no reward. Visual stimuli display, monitoring of eye position, and the synchronization of stimuli with neurophysiological data were performed with in-house software (28) implemented on the MATLAB environment (Mathworks, Natick, MA).

### Surgery and neurophysiology

Once the animals had reached asymptotic performance in the behavioral tasks, a 20-mm diameter recording cylinder was implanted over the prefrontal cortex of each animal. Localization of the recording cylinder and of electrode penetrations within the cylinder was based on MR imaging, processed with the BrainSight system (Rogue Research, Montreal, Canada). Recordings were collected with epoxylite-coated Tungsten electrodes with a diameter of 250 μm and an impedance of 4 MΩ at 1 KHz (FHC Bowdoin, ME). Electrical signals recorded from the brain were amplified, band-pass filtered between 500 and 8 kHz, and stored through a modular data acquisition system at 25 μs resolution (APM system, FHC, Bowdoin, ME). Recordings were obtained and analyzed from areas 8a and 46 of the dorsolateral prefrontal cortex. After reaching adulthood, determined with the developmental indices described above, the animals were again tested in the same tasks that they were originally trained. A new phase of recordings was then performed from the same areas using identical recording methods.

### Neural Data Analysis

Recorded spike waveforms were sorted into separate units using a semi-automated cluster analysis method based on the KlustaKwik algorithm. The firing rate of units was then determined by averaging spikes in each task epoch. In the ODR task, we identified neurons with significant elevation of firing rate in the 500 ms presentation of the cue, the 1500 ms delay period, and the 250 ms response epoch, after the offset of the fixation point. Firing rate in this period was compared to the 1 s baseline fixation period, prior to the presentation of the cue, and neurons with significant difference in firing rate were identified (paired t-test, p<0.05). The location of the receptive field was determined based on responses in the ODR task. Neurons that did not respond with an elevation of firing rate during the ODR task, and for which the receptive field could not be determined, were excluded from analysis.

Population peri-stimulus time histograms (PSTHs) were constructed averaging responses of multiple neurons. Statistical comparisons in the ODR task involved firing rates distributions recorded during the pre-cue fixation, cue, and delay periods of the task. We employed a permutation test using resampling to test the null hypothesis that firing rates of neurons at the young and adult stages came from the same distribution. We randomly reassigned the group labels (young or adult) in 1000 iterations in each test and compared the observed difference to this distribution of permuted differences. We used a three-way ANOVA to compare the influence of factors young/adult stage, cue location, and depth group on the neuron’s firing rate and spatial tuning.

Receiver Operating Characteristic (ROC) analysis was performed, comparing the distribution of responses to the best location and the location diametric to it. The area under the ROC curve represents the probability that an ideal observer can discriminate between a stimulus appearing in the overall best and diametric location firing rate in each trial, based on the relative difference in firing rate between the two stimulus conditions. The analysis was performed for spikes recorded during the entire cue period and delay period, and in a time-resolved fashion, in sliding 50 ms bins, stepped every 50□ms. Statistical analyses of neural data were performed in MATLAB (2022b) and Python 3.8.

## RESULTS

Four male macaque monkeys (*Macaca mulatta*) were initially trained to perform the ODR task (Fig. 1A) during adolescence. Neurophysiological recordings were obtained from areas 8a and 46 of the dorsolateral prefrontal cortex after the onset of puberty (which we refer to as the “young” stage) and in adulthood (“adult” stage). The time of puberty was determined based on morphometric, radiographic, and hormonal measures as documented previously (21). One round of neurophysiological recordings was obtained after the onset of puberty, at a median age of 4.3 years (range: 4.0 – 5.2 years). This experimental stage lasted 3 – 6 months. Monkeys were then returned to their colony and received no further training or exposure to any behavioral task for approximately 1 year. The monkeys were briefly re-introduced to the task and a second round of recordings was obtained. The median age of animals at the onset of the second stage of experiments was 6.3 years (range of ages: 5.6 – 7.3; range of intervals from young stage: 1.6 – 2.1 years).

### Distribution of recordings

A total of 607 neurons were recorded from areas 8a and 46 of the dorsolateral prefrontal cortex in the young stage, and 830 neurons were recorded in the adult stage from the same monkeys. We identified 331 neurons in the young stage and 320 neurons in the adult stage that responded significantly to at least one visual stimulus during the ODR task compared with baseline activity (evaluated with a paired t-test at the P<0.05 level) and selected these neurons for further analysis.

For each electrode penetration, we noted the position where neural activity was first identified (top of the cortex). The depth of each isolated neuron was recorded relative to the top of the cortex, defined in this fashion. We grouped neurons into three groups depending on the depth they were recorded: superficial (0-800 μm), middle (800-1200 μm), and deep (> 1200 μm). Of task-responsive neurons, 191 neurons at the young stage and 210 neurons at adult stage were classified as superficial; 71 neurons at the young stage and 54 neurons at the adult stage were classified as recorded at middle depths; and 47 neurons at young stage and 56 neurons at the adult stage were classified as deep. As this distribution suggests, the superficial layers tended to be oversampled. However, there was no significant difference in mean depth between the young and adult stage (two-tailed t-test, t_631_=1.496, p=0.14, Fig. 2A-B).

**Figure 2.**
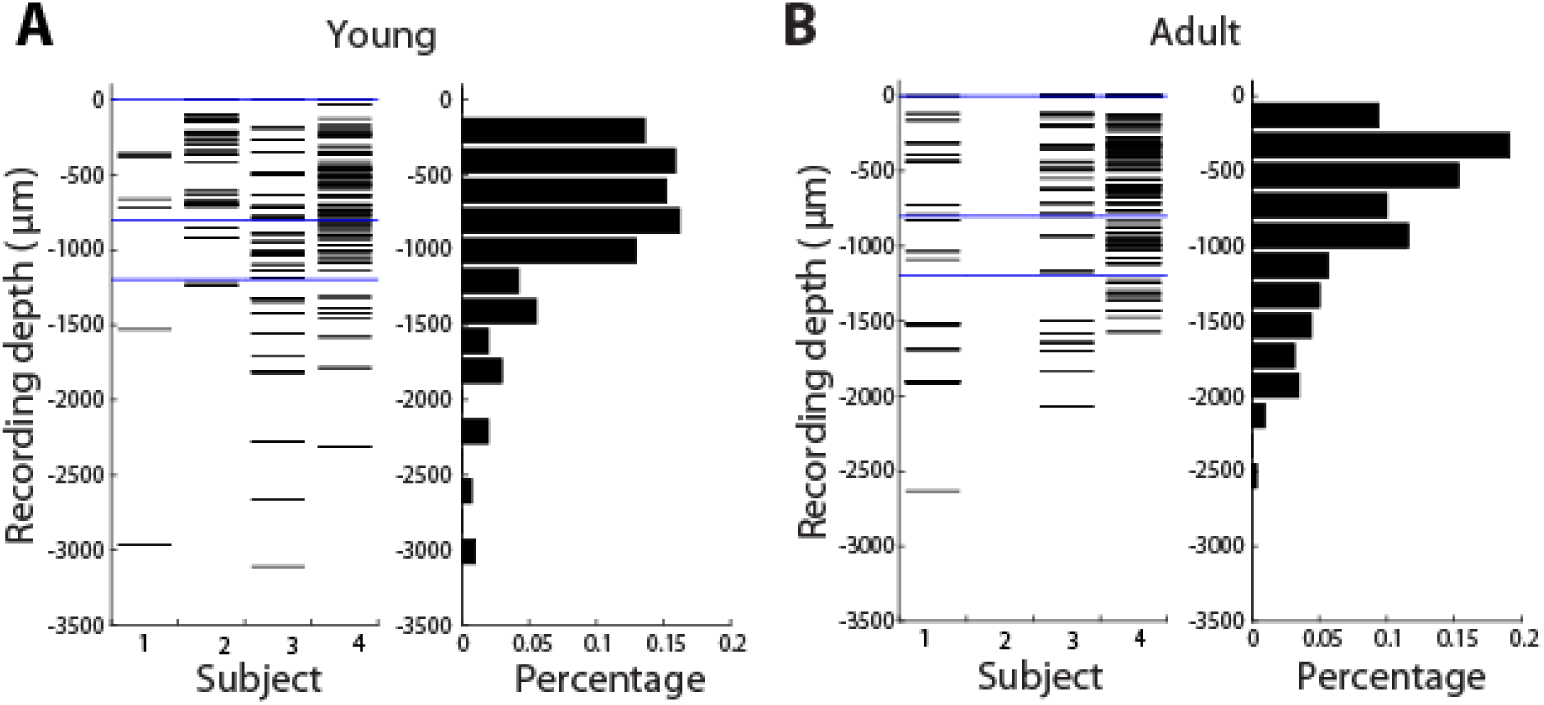
Recording depths. Depths of neurons with significant responses during the task relative to the top of the cortex are shown for the young (A) and adult stage (B). Each horizontal line represents the depth of one neuron. Histograms summarize the depths of neurons in four monkeys in the young and adult stage. Blue horizontal lines indicate the 0, 800 and 1200 μm depths which defined the boundaries for the superficial, middle, and deep groups.

### Firing rate

As we have reported previously, the overall effect of developmental maturation was an increase in firing rate, including for the baseline fixation period, but also for the delay period relative to this baseline (21). We therefore sought to determine whether this increase was confined to superficial layers. A significant increase in pre-cue baseline activity was evident at the adult stage relative to the young stage in the superficial group (mean discharge rate in young stage: 6.50 spikes/s, in adult: 9.30 spikes/s, permutation test, P<0.001). No significant difference in baseline activity was evident in the middle depth group, though a trend toward an increase was present (mean discharge rate: young stage: 8.35 spikes/s, in adult: 12.0 spikes/s, permutation test, P>0.05). Much less of an increase, and no significant difference was present in the deep layer (mean discharge rate: young stage: 8.0 spikes/s, in adult: 8.15 spikes/s, permutation test, P>0.05).

Firing rates during the cue period increased slightly between the adolescent and adult stage, but the difference did not reach statistical significance. The effect was similar for all depth groups. For the superficial depth group, the mean discharge rate increased from 18.55 spikes/s, in the young to 21.3 spikes/s in the adult; in the middle depth group from 21.2 spikes/s in the young and 26.5 spikes/s in the adult; and in the deep group from 19.7 spikes/s in the young to 21.4 spikes/s in the adult (permutation test, P>0.05 for all depth groups). The mean cue period rate relative to the baseline fixation (evoked rate) between stages was not significantly different in any depth group either. The evoked cue firing rates in the superficial depth group were 12.05 spikes/s in the young and 12.0 spikes/s in the adult, in the middle depth group were 12.8 spikes/s in the young and 14.6 spikes/s in the adult; and in the deep group were 11.7 spikes/s in the young and 13.3 spikes/s in the adult (permutation test, P>0.05 for all depth groups).

In contrast, we found delay period discharge rate was significantly higher at the adult stage. A significant increase for the firing rate was observed in the superficial depth group: mean discharge rate rose from 10.7 to 16.38 spikes/s (permutation test, P < 0.001). The mean evoked delay period rate similarly rose from 3.8 spikes/s in the young to 6.9 spikes/s in the adult (permutation test, P < 0.001). An even greater absolute increase was evident in the middle depth group (young: 12.5 spikes/s, adult: 20.2 spikes/s, permutation test, P < 0.05). The difference in evoked delay period rate was also significant in middle depth group (young: 3.8 spikes/s; adult: 8.6 spikes/s; permutation test, P<0.05). In contrast, no significant difference was observed in the deep group for either the absolute firing rate (young: 11.9 spikes/s; adult: 13.8 spikes/s; permutation test, P>0.05) or the evoked firing rate (young: 3.8 spikes/s; adult 5.3 spikes/s, permutation test, P>0.05).

### Tuning and selectivity

We next sought to test whether changes in firing rate also translated in improvement in the discriminability between stimuli. To analyze the discriminability between the best location and its opposite, we used a receiver operating characteristic analysis. We compared area-under-the Receiver Operating Characteristic curve values (auROC) across depth groups. Mean auROC values computed during the stimulus presentation period of the ODR task (Fig. 4) were indistinguishable between the young and adult stages in superficial and mid layers (auROC_superficial_ = 0.799 in adult and 0.807 in young; auROC_middle_ = 0.802 in adult and 0.804 in young; permutation test, P >0.05 in both layers). Discriminability increased in the deep group (auROC_deep_ = 0.846 in adult and 0.798 in young; permutation test, P < 0.05). Mean auROC values for the delay period were higher at the adult stage than young stage in all depth groups, with the larger differences observed in deeper layers (auROC_superficial_ = 0.668 in adult and 0.656 in young, permutation test, P >0.05; auROC_mid_ = 0.686 in adult and 0.625 in young, permutation test, P >0.05; auROC_deep_ = 0.737 in adult and 0.649 in young; permutation test, P < 0.05). This resulted from larger percentages of neurons in the adult reaching higher ROC values at each time point of the delay period (Fig. 4).

The changes in ROC values between depth groups and stages did not correspond to a proportional increase in firing rates for the best stimulus locations (Fig. 3, Fig. 4). Thus, we proceeded to analyze the tuning of the units in different layers, both during the cue presentation and delay period. A three-way ANOVA of firing rate, with factors of cue location, layer, and stage, revealed a significant main effect of depth group, for both the cue presentation period and for the delay period (F_2, 4998_ = 19.18, P = 5.02 x 10^-9^ for the cue period; F_2, 4998_ = 21.77, P = 3.84 x 10^-10^ for the delay period). Similarly, the test revealed a significant main effect of stage, and this was present both in the cue period and in the delay period (F_1, 4998_ = 16.43, P = 5.12 x 10^-5^ for the cue period; F_1, 4998_ = 147.64, P = 1.68 x 10^-33^ for the delay period). A significant interaction between group and stage was also present (F_2, 4998_ = 6.26, P =0.0019 for the cue period; F_2, 4998_ = 17.99, P = 1.64 x 10^-8^ for the delay period). These results suggest significant differences in firing rate between depth groups, and differential effects of developmental stage on firing rate. On the other hand, there was no significant interaction between cue location and depth group (F_14, 4998_ = 0.31, P = 0.990 for the cue period; F_14, 4998_ = 0.22, P = 0.998 for the delay period). The result suggests that the shape of tuning curves was similar in different depth groups.

**Figure 3.**
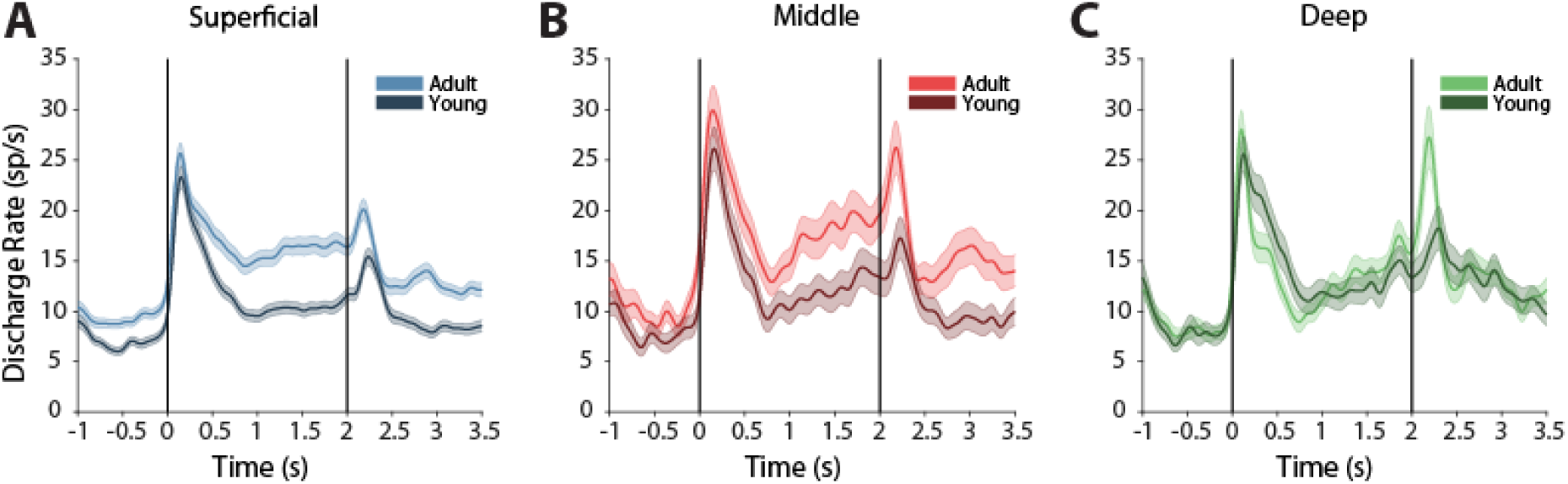
Firing rate in different depth groups and developmental stages. A. Average, population peri-stimulus time histogram for neurons that responded to the visual stimulus and were recorded during the ODR task from superficial layer at adult and young stages. Responses are shown for a stimulus in the neuron’s receptive field and aligned to the stimulus onset of each trial. Vertical lines represent stimulus onset and fixation offset. B. As in A, for the mid layer. C. As in A, for the deep layer.

**Figure 4.**
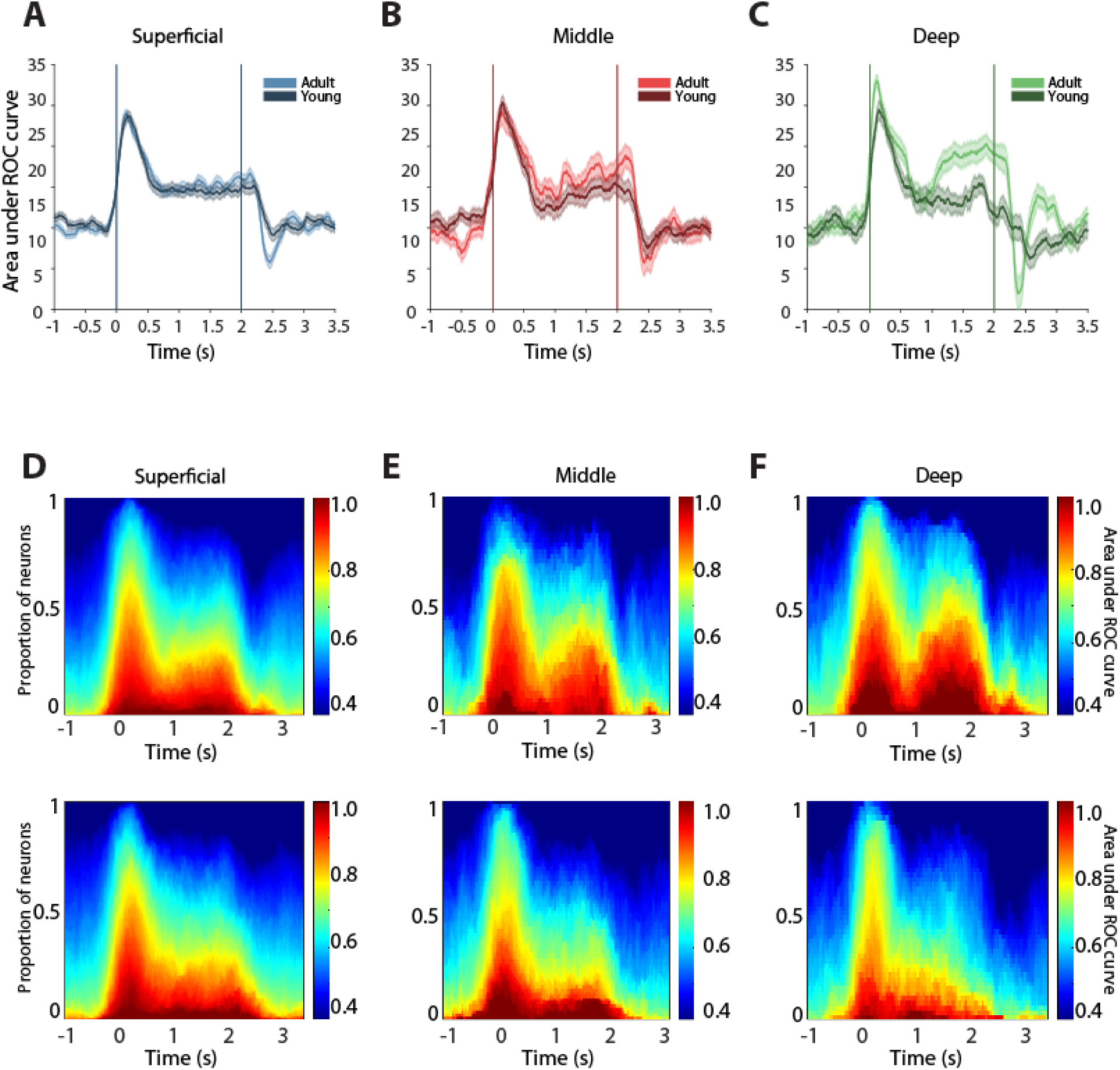
ROC analysis in each layer. A. Area under ROC curve in successive 100□ms windows is plotted as a function of time during the ODR task, for superficial layer neurons at adult and young stages. B, C. As in A, for the mid layer and deep layer, respectively. D. Percentages of neurons at adult (top) and young (bottom) stages reaching different levels of ROC values at each time point of the ODR task. E, F. As in D, for the mid layer and deep layer, respectively.

The best cue location exhibited the greatest increase in firing rates in all layers, and the relative difference was more pronounced in the delay than in the cue period (Fig. 5a, b). Unlike in superficial and mid layers, firing rates of locations away from the neurons’ best locations were lower in deep layer (Fig. 5), although the difference did not reach statistical significance. A two-way ANOVA was performed on each layer individually. We observed significant effects of the stage on cue firing rate and delay activity in superficial and mid layers with greater increases in the superficial depth group (F_1, 3206_ = 14.99, P = 0.0001 for the cue period, F_1, 3206_ = 150.87, P = 6.39 x 10^-34^ for the delay period in superficial group; F_1, 998_ = 7.21, P = 0.007 for the cue period, F_1, 998_ = 29.22, P = 8.06 x 10^-8^ for the delay period in the middle depth group), and no significant effect of stage in the deep group (F_1, 822_ = 1.69, P = 0.194 for the cue period; F_1, 822_ = 0.31, P = 0.575 for the delay period).

**Figure 5.**
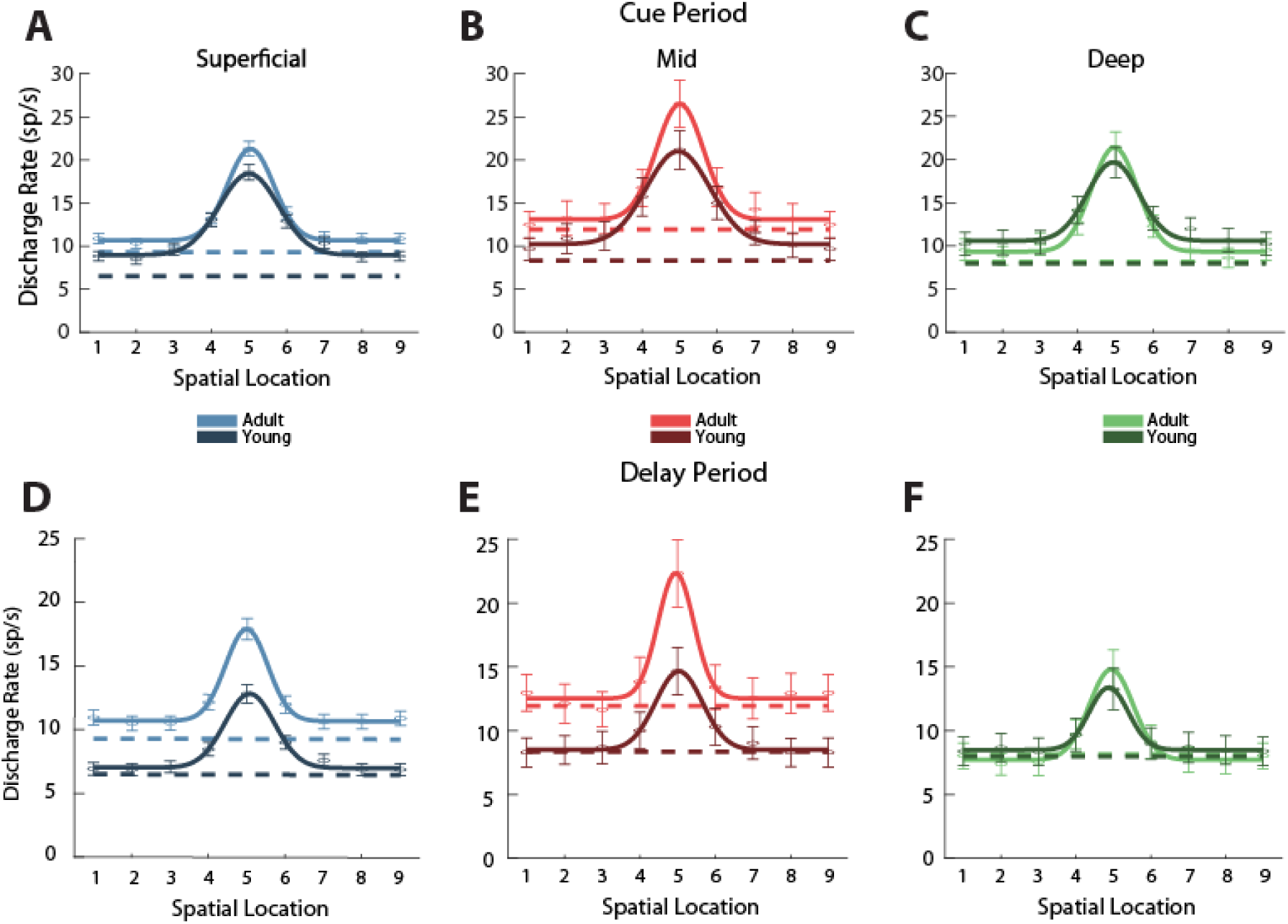
Neuronal tuning in each layer. A. Average activity (and s.e.m.) during the cue period of the ODR task in neurons recorded from superficial layer at adult and young stages. Locations have been rotated, so that the best location of each neuron is represented in location 5. Location 9 is the same as location 1. Solid lines represent the best Gaussian fit of the population average. B, C. As in A, for the mid layer and deep layer, respectively. D. Average activity (and s.e.m.) during the delay period of the ODR task from superficial layer at adult and young stages. E, F. As in D, for the mid layer and deep layer, respectively.

## DISCUSSION

Changes in the activity of the prefrontal cortex have been thought to underlie maturation of cognitive abilities during adolescence, including working memory (29, 30) and other executive functions (31, 32). These in turn are associated with structural brain changes, including changes in cortical volume and thickness that occur during this period (11, 33). Non-human primate species undergo a similar pattern of structural brain changes (34, 35). Neurophysiological studies in non-human primates have begun to uncover the nature of these changes at the level of single neuron responses (21, 22). Our current study provides a comprehensive analysis of the developmental changes in the firing rates and selectivity of neurons in the dorsolateral prefrontal cortex of macaque monkeys at different cortical depths. We found that increases in firing rate between adolescence and adulthood were not uniform across all layers of the cortex, but changes were not confined to the superficial layers, either, which are most commonly thought to mediate working memory responses (23, 25, 26). Our findings reveal a complex picture of neural maturation in the prefrontal cortex during adolescence, characterized by layer-specific changes in firing rate and stimulus discriminability. This layer-specific maturation pattern suggests that different cortical layers may play different roles in working memory processing, which mature differently over the course of development.

### Superficial layers in working memory maintenance

Horizontal connections between pyramidal neurons in layers II and III of the prefrontal cortex have been thought to provide the anatomical substrate through which neuronal discharges reverberate and persist during working memory (23, 36). Biophysical models of working memory have simulated precisely this type of connections between neurons to create recurrent networks of units that continue to excite each other (37, 38). Indeed, neurons with persistent activity are more readily identified in the superficial layers of the prefrontal cortex, comprising the top ∼800 μm of the cortical volume (25). Other types of neural activity that has been associated with working memory, such as synchronized gamma-band oscillations in the local field potential, have also been localized in the superficial layers (39). Recent advancements in layer-specific functional MRI techniques have also allowed for the dissociation of activity in superficial and deeper cortical layers during different periods of a working memory task in the human dorsolateral prefrontal cortex (40).

Given this evidence, it appeared plausible that improvement in working memory ability would manifest itself with improved persistent activity generation, focused on the superficial layers. A systematic difference in firing rate between the adolescent and adult stage was indeed present for the superficial layers; however, this involved an overall increase in fixation-period rather than delay-period activity. This “baseline” activity, rather than being passively generated, has been increasingly recognized as a preparatory process that has important implication for the execution of cognitive task (41, 42). The generation of persistent activity during working memory is directly dependent on this baseline; computational studies reveal that networks with high baseline activity are more likely to be able to sustain activation; persistent activity may die off in networks with lower baseline activity if neurons fail to generate sufficient action potentials as a result of synaptic activation (43). In that sense, the increased baseline activity we observed in the superficial layers of the adult prefrontal cortex may be indirectly related to the improved working memory ability. It was also notable that this baseline firing rate increase was less pronounced as a function of cortical depth; the middle depth group showed only a slight, non-statistically significant increase, whereas the deep group showed the least change, with the mean discharge rate remaining almost the same from the young to the adult stage.

### Delay period activity

The biggest increase in delay period activity relative to baseline was observed for the middle layers; superficial layers exhibited a more modest increase in this measure. This result was unexpected, as middle layers have been more implicated in the representation of sensory stimuli (25), driven by afferent inputs to the prefrontal cortex. The result raises the possibility that much of the increase in delay period activity is a distributed effect, contributed by areas connected to the prefrontal cortex (44), as much as it is dependent on intrinsic changes within the prefrontal cortex.

There are several caveats in this analysis. Layer determination was only approximate, with three depth ranges identified based on fixed depths, rather than precise histological identification. Depth was also determined approximately, based at the point that spiking was first detected. However, it should be noted that this measure of depth was conservative; it is much more likely for the true top of the cortex to be missed during a penetration than falsely be identified prematurely. In this sense, our results may even underestimate the adolescent changes in non-superficial layers.

Another caveat is that adult monkeys had more cumulative exposure to the task than young monkeys, and it is well established that training can increase persistent activity during working memory (45, 46). In this sense, the changes we observed in the middle layers may be primarily driven by training rather than by developmental maturation. We have shown changes in firing rate cannot be fully attributed by training or improved performance attained in the task (21). Nonetheless, it would be no less surprising that effects of training would disproportionately affect the middle layers rather than the superficial ones during the generation of persistent activity.

### Stimulus representation and discriminability

In yet another unexpected finding, the increase in delay firing rate did not lead to a proportional improvement in stimulus discriminability in the superficial and middle layers. In contrast, it was the deep layers that exhibited the greatest improvement in stimulus discriminability from the young to the adult stage despite the lack of a significant increase in delay firing rate. Deep layers are thought to reflect the output of the maintenance operation for the guidance of motor action (25), and in this sense, increased precision may be most critical there. This layer-specific improvement in discriminability may reflect a refinement of neural circuits in the deep layer, which enhances the precision of neural tuning and contributes to the maturation of cognitive abilities. In fact, evidence exists that a multitude of signals is communicated from frontal cortical areas to subcortical structures such as the superior colliculus during the maintenance of working memory (47). Thalamic nuclei, and particularly the mediodorsal nucleus also receive input from deep cortical layers and exhibit persistent activity during the delay period of working memory tasks, whose disruption impairs performance (48, 49).

Our observations of layer-specific changes in firing rate and stimulus discriminability underscore the importance of considering the structure–function coupling at a fine spatial scale. Previous studies have uncovered laminar fractionation of other cognitive functions as well (50, 51). Future studies should consider the heterogeneity of neural development across different cortical layers and aim to elucidate the underlying mechanisms and functional implications of these layer-specific changes.

## ACKNOWLEDGEMENTS

We thank Stanley Vinet and Chrissy Suell for technical assistance and Rye Jaffe for helpful comments on the manuscript. Research reported in this paper was supported by the National Institute of Mental Health under award number R01 MH116675.

